# An efficient method for immortalizing mouse embryonic fibroblasts

**DOI:** 10.1101/2024.06.27.601088

**Authors:** Srisathya Srinivasan, Hsin-Yi Henry Ho

## Abstract

Mouse embryonic fibroblasts (MEFs) derived from genetically modified mice are a valuable resource for studying gene function and regulation. The MEF system can also be combined with rescue studies to characterize the function of mutant genes/proteins, such as disease-causing variants. However, primary MEFs undergo senescence soon after isolation and passaging, making long-term genetic manipulations difficult. Previously described methods for MEF immortalization are often inefficient or alter the physiological properties of the cells. Here, we describe an optimized protocol for immortalizing MEFs via CRISPR-mediated deletion of the *Tp53* gene. This method is highly efficient and consistently generates immortalized MEFs, or iMEFs, within 14 days. Importantly, iMEFs closely resemble the parent cell populations, and individual iMEFs can be cloned and expanded for subsequent genetic manipulation and characterization. We envision that this protocol can be adopted to immortalize other mouse primary cell types.

**Key Features:**

- CRISPR-based knockout of the *Tp53* gene enables efficient immortalization of mouse embryonic fibroblasts (MEFs) in under 2 weeks.
- Immortalization requires a Neon electroporator or another comparable electroporation system to transfect cells with the *Tp53* CRISPR constructs.

## Background

The ability to culture cells long-term *in vitro* has been a cornerstone of modern cell biological research. Except for stem and cancer cells, most primary cells undergo senescence in culture, making immortalization an essential step during the establishment of permanent cell lines from native tissues. Earlier methods of cell immortalization involved overexpression of oncogenes to transform cells (Bryan & Reddel, 1994; Graham, Smiley, Russell, & Nairn, 1977; Hahn et al., 2002; Newbold, Overell, & Connell, 1982). Though efficient, these methods frequently result in the acquisition of cancer-like phenotypes, including the loss of contact inhibition and anchorage dependent growth, as well as alterations in growth factor requirement, metabolism and other signaling activities (Kelekar & Cole, 1987; Pipas, 2009; Price, Moorwood, James, Burke, & Mayne, 1994).

Overexpression of telomerase (TERT, or <u>te</u>lomerase <u>r</u>everse transcriptase) is another common method to transform mammalian cells (Bodnar et al., 1998). This method is highly effective in immortalizing human cells and has a lower tendency to induce cancer-like phenotypes (Wieser et al., 2008). However, few examples exist in the literature demonstrating that TERT overexpression alone can immortalize mouse cells (Steele et al., 2010), suggesting that this method is generally less effective toward mouse cells. In our own experience, we have not been able to successfully immortalize mouse embryonic fibroblasts via overexpression of telomerase.

3T3 cells are widely used mouse embryonic fibroblast (MEF) lines spontaneously immortalized through serial passaging. The ‘3T3’ method, originally described by Howard Green and colleagues, involves seeding primary MEFs at <u>3</u> × 10^5^ cells/plate and transferring every <u>3</u> days at the same density until rapidly dividing, immortalized cells emerge (Todaro & Green, 1963). While this immortalization process is considered gentle and does not cause neoplastic transformation (Todaro & Green, 1963), it is inefficient and time consuming. Moreover, because the method relies on mutation(s) that spontaneously arise during the prolonged cell passaging phase, it is difficult to compare cell lines derived from different immortalization experiments.

Importantly, Harvey and Levine (Harvey & Levine, 1991) reported that MEF lines immortalized via the 3T3 method frequently carry loss-of-function mutations in the tumor suppressor gene *Tp53*, suggesting that this is a key immortalizing event. Consistent with this observation, we found that CRISPR/Cas9-mediated ablation of the *Tp53* gene robustly immortalizes primary MEFs. We have used the protocol described here to generate immortalized MEF lines, or iMEFs, in under two weeks and from as few as 50,000 primary cells. iMEF lines have also been successfully established from WT and genetically modified embryos of different genetic backgrounds (C57BL/6 and mixed C57BL/6; 129J). Lastly, iMEFs generated through *Tp53* deletion can be subcloned and genetically manipulated further in downstream applications, such as gene rescue experiments.

## Materials and Reagents

### Biological Materials

1. Pregnant mice carrying embryonic day 12.5 (E12.5) embryos.

### Reagents

1. Phosphate buffered saline (PBS) without Ca^2+^ and Mg^2+^ (Cytiva, catalog number: SH30256.01)
2. Hanks’ balanced salt solution (HBSS) without Ca^2+^ and Mg^2+^ (Gibco, catalog number: 14170112)
3. Trypsin from bovine pancreas (Worthington Biochemical, catalog number: LS003702) – for MEF preparation.
4. RQ1 RNase-free DNase (Promega, catalog number: M6101)
5. Non-essential amino acids solution in minimal essential medium (MEM NEAA) (100X, Gibco, catalog number: 11140050)
6. Sodium pyruvate (100 mM, Gibco, catalog number: 11360070)
7. Dulbecco’s modified Eagle’s medium, high glucose (4.5g/L) (DMEM) (1X, Gibco, catalog number: 11960-044)
8. Fetal bovine serum (FBS), United States qualified grade (Gibco, catalog number: 26-140-079)
9. Penicillin-streptomycin (100X, Gibco, catalog number: 15-140-122)
10. L-glutamine (200 mM, Gibco, catalog number: 25030081)
11. Dimethyl sulfoxide (DMSO), sterile, tissue culture grade (Millipore Sigma, catalog number: D2650-100ML)
12. 0.25% trypsin-EDTA with phenol red (Gibco, catalog number: 25200072) – for cell harvest and passaging after the initial MEF preparation.
13. Px461-Cas9n-Trp53-sgRNA-alpha plasmid (Addgene, plasmid number: 88846)
14. Px461-Cas9n-Trp53-sgRNA-beta plasmid (Addgene, plasmid number: 88847)
15. pCAG-GFP plasmid (Addgene, plasmid number: 11150)

### Solutions

1. 0.1% trypsin for MEF prep (See Recipes)
2. Complete cell culture media (culture media) (See Recipes).
3. Cell freezing media (See Recipes)

### Recipes

#### 1. 0.1% trypsin for MEF preparation

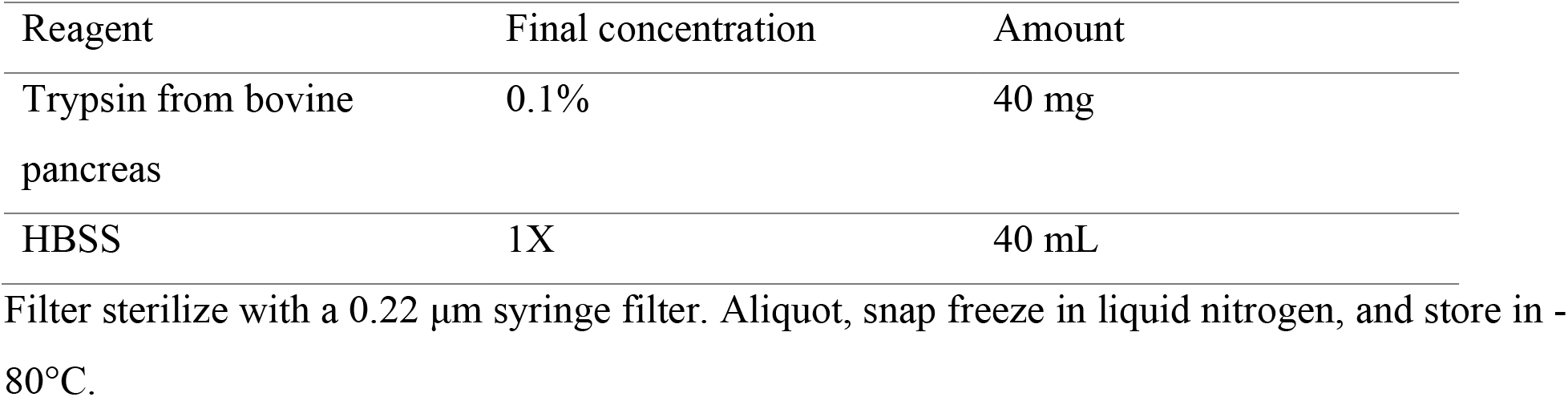

#### 2. Complete cell culture media (culture media)

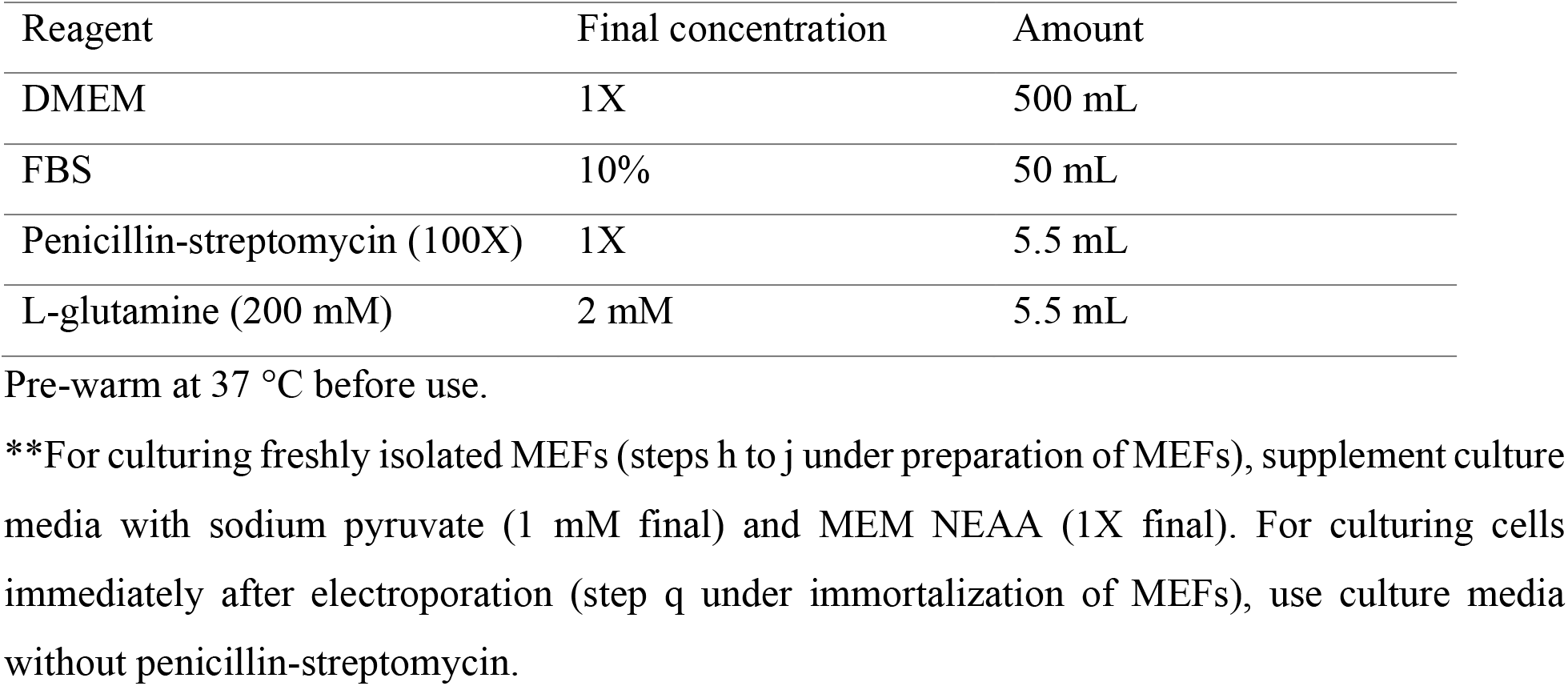

Filter sterilize with a 0.22 μm syringe filter. Aliquot, snap freeze in liquid nitrogen, and store in -80°C. 2. Complete cell culture media (culture media)
| Reagent | Final concentration | Amount |
| --- | --- | --- |
| DMEM | 1X | 500 mL |
| FBS | 10% | 50 mL |
| Penicillin-streptomycin (100X) | 1X | 5.5 mL |
| L-glutamine (200 mM) | 2 mM | 5.5 mL |

Pre-warm at 37 °C before use.
**\*\***For culturing freshly isolated MEFs (steps h to j under preparation of MEFs), supplement culture media with sodium pyruvate (1 mM final) and MEM NEAA (1X final). For culturing cells immediately after electroporation (step q under immortalization of MEFs), use culture media without penicillin-streptomycin.

#### 3. Cell freezing media

| Reagent | Final concentration | Amount |
| --- | --- | --- |
| Culture media | 90% | 45 mL |
| DMSO | 10% | 5 mL |

Filter sterilize with a 0.22 μm syringe filter.

### Laboratory Supplies

1. Surgical scissors (Fine Science Tools, model: Surgical scissors-sharp, catalog number: 14002-12)
2. Adson forceps (Fine Science Tools, model: Adson forceps, catalog number: 11006-12)
3. Fine tip forceps (Fine Science Tools, model: Dumont #5 Inox, catalog number: 11251-20)
4. 60 mm TC-treated cell culture dishes (Corning, Brand: Falcon, catalog number: 353002)
5. 100 mm TC-treated cell culture dishes (Corning, brand: Falcon, catalog number: 353003)
6. 6-well clear TC-treated multi well plates (Corning, Brand: Costar, catalog number: 3516)
7. 24-well clear TC-treated multi well plates (Corning, Brand: Costar, catalog number: 3524)
8. 50 mL centrifuge tubes (Thermo Fisher Scientific, brand: Nunc, catalog number: 339653)
9. Pipettes (Eppendorf: Research Plus, P-1000, P-200, P-20, catalog number: 3123000918)
10. Filtered pipette tips (Fisher Scientific, brand: SureOne, catalog numbers: 02-707-442; 0.1 to 10 μL, 02-707-432; 2 to 20 μL, 02-707-430; 20 to 200 μL, 02-707-404; 100 to 1000 μL)
11. Metal tube rack for 1.5 mL tubes (Stratagene, catalog number: 41018. The item has been discontinued, but any similar rack will work)
12. 5 mL serological pipettes (Thermo Fisher Scientific, brand: Nunc, catalog number: 170355N)
13. Pipette controller (Drummond, brand: Pipet-Aid, catalog number: 4-000-100)
14. 1.5 mL microcentrifuge tubes (Denville Scientific, brand: PosiClick, catalog number: C2170)
15. Neon transfection system 10 μL kit (Thermo Fisher Scientific, brand: Neon, catalog number: MPK1096)
16. Cell strainers (Corning, brand: Falcon, catalog number: 352340)
17. 30 mL syringes (BD, catalog number: 302832)
18. 0.22 μm syringe filters (Thermo Fisher Scientific, brand: Fisherbrand, catalog number: 09-719C)

### Equipment

1. Dissecting microscope (Leica, model: S7E, catalog number: S7E-PS)
2. Biological safety cabinet (Thermo Fisher Scientific, model: 1300 series class II, catalog number: 1323TS)
3. CO_2_ incubator (Thermo Fisher Scientific, model: Heracell 150i, catalog number: 51026281), set at 37 °C and 5% CO_2_.
4. Benchtop centrifuge (Thermo Fisher Scientific, model: Sorvall ST 8, catalog number: 75007200)
5. 8 × 50 swing bucket rotor (Thermo Fisher Scientific, model: TX-100S, catalog number: 75005704)
6. Microcentrifuge (Thermo Fisher Scientific, model: Sorvall Legend Micro 21, catalog number: 75002436)
7. 24 × 1.5/2.0mL rotor with ClickSeal Lid (Thermo Fisher Scientific, catalog number: 75003424)
8. Ultra-low (−80°C) freezer (Thermo Fisher Scientific, model: Revco UxF, catalog number: UXF60086A)
9. Liquid nitrogen storage unit (Thermo Fisher Scientific, model: Cryoplus 3, catalog number: 7404)
10. Cell counter (Corning, catalog number: 6749)
11. Electroporation device (Thermo Fisher Scientific, model: Neon, catalog number: MPK5000) – this instrument is discontinued. Other similar electroporation devices can be used, but optimization will be required.
12. Inverted fluorescence microscope (Thermo Fisher Scientific, model: Evos FL, catalog number: AMF4300)
13. Mr. Frosty freezing container (Thermo Fisher Scientific, catalog number: 5100-0001)

### Procedure

#### A. Preparation of primary mouse embryonic fibroblasts (MEFs)

*Performing the embryo dissection in a laminar flow hood is ideal. However, if a laminar flow hood is not available, the procedure can be performed on a standard lab bench. All solution stocks (e*.*g. HBSS) should be kept sterile and transferred to the dishes used for dissection in a biosafety cabinet. General procedures on mouse husbandry can be found in (Silver, 1995)*.

##### 1. Dissection of E12.5 embryos from timed mating

a. Set up timed mating and start checking the females for vaginal plugs the following morning. *Critical: Plugged females should be separated from the male and housed until 12*.*5 days post-coitum*.
b. Euthanize the female at 12.5 days post-coitum via CO_2_ inhalation, followed by cervical dislocation to ensure death.
c. Spray the mouse with 70% ethanol to sterilize the abdominal surface.
d. Dissect out the uterine horns and rinse in PBS in 100 mm dishes. *Critical: When dissecting the uterine horns, ensure that the intestines are not nicked, and the uterine horns do not touch the fur to prevent contamination*.
e. Separate each embryo from the uterus and place it a 60 mm dish containing HBSS.
f. In the same 60mm dish and under a dissecting scope, remove the placenta and embryonic sac. *Notes:*
  i. *The embryonic sacs can be collected and used to genotype the embryos*.
  ii. *The sac should be gripped firmly with fine tip forceps and rinsed briefly under a slow stream of running DI water before transferring to a collection tube. This step helps to prevent cross-contamination from maternal tissues*.
g. Remove the eyes, internal organs, and the brain of the embryo as much as possible using fine tip forceps and discard.
h. Transfer the dissected embryo to a 6-well plate filled with HBSS and keep the plate on ice until all embryos have been dissected. *Critical: The plate(s) is kept on ice to ensure the viability of the embryonic tissues. Proceed to the next step (preparation of MEFs) as soon as possible*.

##### 2. Preparation of MEFs

*This and all subsequent steps should be performed in a biosafety cabinet under sterile conditions*.

*Before starting:*

- *Pre-warm the metal rack at 37* °*C in the TC incubator*.
- *Thaw 0*.*1% trypsin (from bovine pancreas) and keep it on ice once fully thawed*.
- *Add 0*.*5 mL of 0*.*1% trypsin and 50 μL of Promega DNaseI to a sterile 1*.*5 mL tube. Mix well with a P1000 pipette. Prepare as many tubes as there are embryos to process*.

a. Transfer the first embryo from the 6-well with HBSS into a 1.5 mL microcentrifuge tube with trypsin and DNaseI using fine tip forceps.
b. Homogenize the embryo by pipetting up and down 12 to 15 times using a P1000 pipette, or until embryo is broken into very small chunks. Store the tube with the homogenized embryo on ice and proceed to homogenize the next embryo (steps a to b), until all embryos are homogenized. *Critical: It is very important to not over-homogenize the embryo as this can cause cell lysis. When the homogenate starts to feel increasingly viscous, it is over-homogenized*.
c. Transfer the tubes with the homogenized embryos to the pre-warmed metal rack and incubate at 37 °C for 8 minutes exactly.
d. While the incubation is ongoing, add 5 mL of cold culture media to a 50 mL conical tube in preparation for the next step. Prepare as many tubes as there are embryos. *Note: The media should be cold to facilitate neutralization of the trypsin*.
e. After the 8-minute incubation at 37 °C, transfer the homogenates to the 50 mL conical tubes with cold culture media (prepared in step d) using a P1000 pipette.
f. Use a 5 mL serological pipette to pipette the mixture up and down 8 to 10 times very gently.
g. Centrifuge the neutralized homogenates at 500 x g for 10 minutes at room temperature in a centrifuge equipped with a swing bucket rotor. If a refrigerated centrifuge is available, centrifuge at 4 °C.
h. While centrifugation is ongoing, add 9 mL of pre-warmed culture media supplemented with 100 μL sodium pyruvate (1 mM final) and 100 μL MEM NEAA (1X final concentration) to a 100 mm TC-treated cell culture dish. Prepare as many dishes as there are embryos. Keep the dishes in the TC incubator.
i. Once the centrifugation is done, carefully remove the supernatant by aspiration. *Note: Be careful to avoid disturbing the pellet or getting the aspirator tip too close to the pellet, as the pellet can easily get aspirated away*.
j. Gently resuspend each pellet in 1 mL culture media using a P1000 pipette and transfer the cell suspension to the pre-warmed 100 mm dish with 9 mL MEF media (prepared in step h).
k. Shake the plate gently to ensure uniform distribution of the cells.
l. Incubate the cells in a TC incubator (37 °C, 5% CO_2_) without disturbance until the next day.
m. Next morning, observe the cells to assess cell health. *Note: The MEFs should be approximately 20 to 40% confluent 1 day after isolation and get fully confluent within 2 to 3 days. Some cell death is expected, but the majority of cells should survive and adhere to the culture dish by the morning after isolation (Figure 1)*.
n. Once the cells are confluent, wash once with PBS and trypsinize the cells in each 10 mm dish with 3 mL of prewarmed 0.25% trypsin-EDTA.
o. Once the cells lift off the plate (∼1 to 2 minutes), neutralize the trypsin with 7 mL culture media.
p. Filter the cell suspension (from step o, still in the trypsin/culture media mixture) through a 40 μm cell strainer and collect in a 50 mL tube. *Note: This step removes large cell/tissue clumps*.
q. Centrifuge the cell suspension at 500 x g for 5 minutes at room temperature in a centrifuge equipped with a swing bucket rotor.
r. Carefully discard the supernatant by aspiration. At this point, cells can be split and cultured further (step s below) or frozen (step t below).
s. To culture the MEFs further, resuspend the cell pellet in 1 mL culture media and pipette 12 to 15 times with a P1000 pipette. Proceed with splitting into the desired number of dishes (no more than a 1 to 5 split).
t. To freeze the MEFs, resuspend the cell pellet from each 100 mm dish in 1 mL cell freezing media and transfer to a cryovial. Put the cryovials in a room temperature Mr. Frosty freezing container, then store in an ultra-low freezer at -80°C overnight. For long-term storage, transfer cryovials to a liquid nitrogen freezer. *Pause point: Primary MEFs can be stored in liquid nitrogen indefinitely*. *Caution: Precautions (eye protection, cryosafe gloves and lab coat) must be taken while handling liquid nitrogen*.

**Figure 1:**
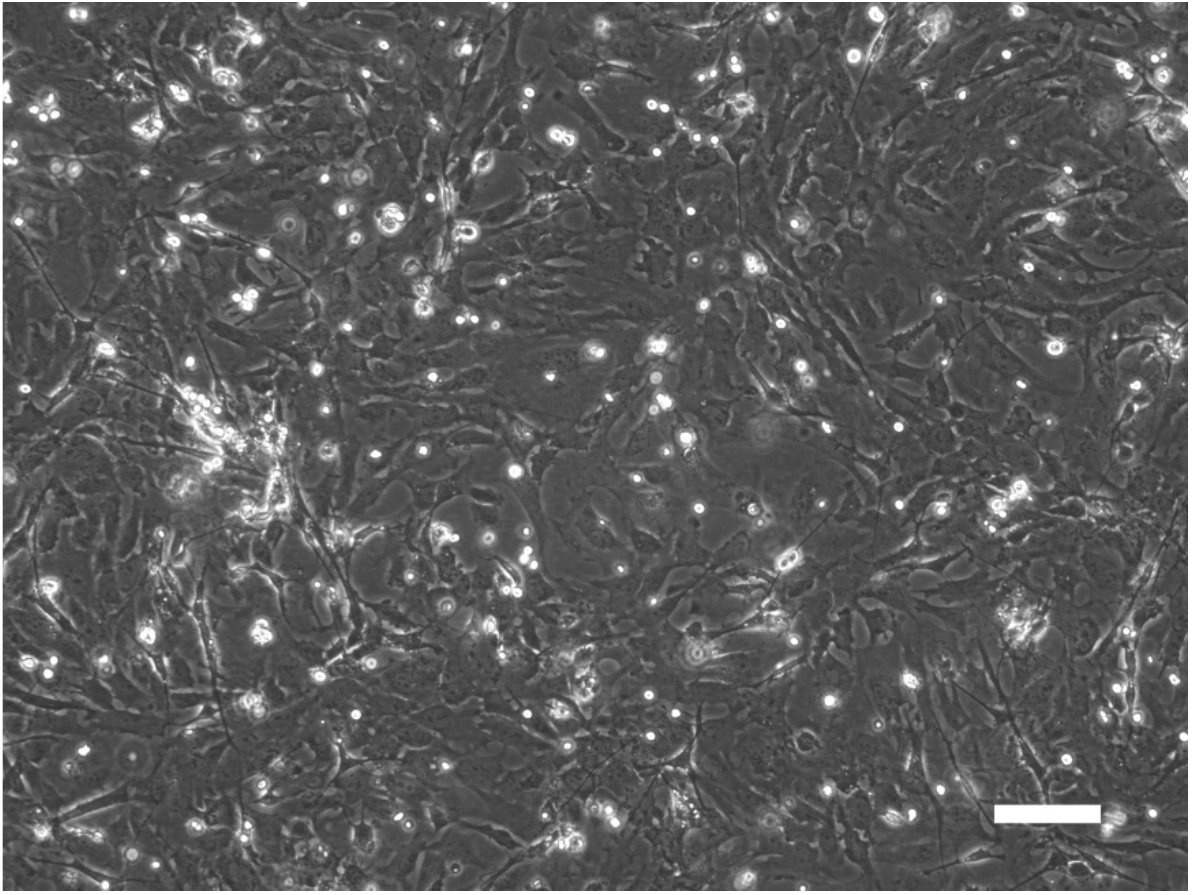
Representative image of MEF from C57BL/6 embryos (E12.5) 48 hours after isolation. Scale bar: 100μm. Magnification: 10X.

#### B. Immortalization of MEFs

*This protocol uses 2 different electroporation programs to maximize the chance of successful transfection and cell immortalization. If both programs are successful, the cells can be pooled*.

##### Before starting

- *∼2 days before electroporation – seed a MEF stock such that cells reach 70 to 90% confluency and are growing robustly on the day of electroporation. Cells can be split from a live culture or thawed from a frozen stock. Cells should be under 3 passages. We routinely use passage 0 (P0) or P1 cultures. A minimum of 400,000 cells are required for the experiment. We typically prepare at least 1 to 2 million cells (two wells of a 6-well plate)*
- *Before starting the electroporation experiment – prepare culture media without penicillin-streptomycin (10 mL) and add 500 μL /well to a 24-well plate; pre-warm at 37°C. Also pre-warm 0*.*25% trypsin-EDTA at 37°C*.
- *Review the Neon electroporator guide (https://assets.thermofisher.com/TFS-Assets/LSG/manuals/neon_device_man.pdf; https://www.thermofisher.com/content/dam/LifeTech/migration/en/filelibrary/cell-culture/neon-protocols.par.71910.file.dat/mouse%20embryonic%20fibroblasts%20(mef)-embryo.pdf).*

a. Wash cells once with PBS (1 mL for each 6-well) and trypinize with 0.25% trypsin-EDTA (0.3 mL for each 6-well).
b. Incubate at 37 °C until the cells lift off the plate (∼1 to 2 minutes).
c. Neutralize the trypsin with culture media without penicillin-streptomycin (0.7 mL for each 6-well) and transfer the cell suspension to a 1.5 mL microcentrifuge tube or another appropriately size tube.
d. Pipette the cells 8 to 10 times using a P1000 pipette to achieve a single cell suspension.
e. Count the cells and determine the cell density (cells are still in the trypsin/culture media mixture at this point).
f. Transfer 400,000 cells into a new 1.5 mL microcentrifuge tube (each transfection requires 50,000 cells; this is therefore enough for 8 transfections; the actual number of transfections to be performed is 6).
g. Centrifuge the cells at 500 x g for 5 minutes in a microcentrifuge.
h. Resuspend the cells in 0.5 mL PBS. Centrifuge to pellet the cells (as in step g). *Critical: This step removes the residual trypsin and culture media and is critical for successful electroporation*.
i. Carefully aspirate the PBS and resuspend the cell pellet in 80 μL of Buffer R from the Neon transfection system kit. This is equivalent to a cell density of 5×10^6^ cells/mL in Buffer R. *Note: Remove as much PBS as possible without disrupting the cell pellet, which is very small, before adding Buffer R*.
j. For each of the GFP control and *Tp53* CRIPSR transfections, prepare a 3X reaction master mix according to the table below. *Note: Each individual electroporation reaction requires 10 μL of the cell suspension. For each of the GFP and Tp53 CRISPR groups, two separate electroporation reactions will be performed, one using electroporation Program A and the other using Program B. To account for pipetting errors, a 3X reaction master mix will be prepared*.

k. Mix the cells and DNA gently by pipetting using a P200 pipette.
l. Fill the Neon tube with 3 mL buffer E and insert into the pipette station of the Neon transfection system. *Note: The pipette station should be placed in the biosafety cabinet to ensure sterility. The pulse generator can be kept outside the biosafety cabinet*.
m. Insert a Neon tip into the Neon pipette by pressing the push-button on the Neon pipette to the second stop. Firmly insert the Neon pipette into the Neon tip. Gently release the push-button while still applying some downward pressure to ensure a tight fit.
n. Pipette 10 μL cell/DNA suspension from the *Tp53* electroporation master mix. *Critical:*
  i. *Ensure that there are no air bubbles trapped in the tip*.
  ii. *Ensure that the cells in the master mix are in a uniform suspension. If cells have settled, mix by pipetting up and down using the Neon pipette*.
o. Insert the Neon pipette into the Neon tube with buffer E.
p. Electroporate using Program A below:

|  | GFP (3X reaction master mix) | *Tp53* CRISPR (3X reaction master mix) |
| --- | --- | --- |
| MEF suspension in Buffer R (density: $5 \times 10^6$ cells/mL) | 30 μL | 30 μL |
| pCAG-GFP (1 μg/μL) | 1.5 μL | -- |
| Px461-Cas9n-Trp53-sgRNA (1 μg/μL) | -- | 0.75 μL |
| Px461-Cas9n-Trp53-sgRNA-beta (1 μg/μL) | -- | 0.75 μL |

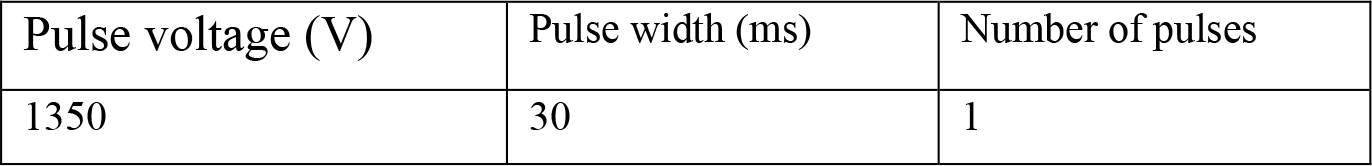 *Note: A very small spark may occur during the pulse. If a large visible spark is seen, it could indicate an issue in the conductance due to a trapped air bubble.*
q. Immediately transfer the cells from the Neon pipette tip into the prepared 24-well plate (pre-warmed culture media without penicillin-streptomycin). Gently shake the 24-well plate to evenly distribute the cells.
r. Repeat *Tp53* electroporation (steps n to q) using Program B below (the same tip can be used):

s. Using the same tip previously used in steps n to r, perform electroporation of the GFP control construct using Programs A and B. *Note: We routinely use each tip for up to 6 electroporations. We electroporate cells with the Tp53 CRISRP constructs first to avoid cross-contaminating them with the GFP control plasmid. However, fresh tip can be used for each electroporation if desired*.
t. Incubate the cells in a TC incubator (37 °C; 5% CO_2_). *Note: Cells can be observed for viability ∼2 hours after electroporation. Critical: Plate disturbance should be kept to a minimum*.
u. Evaluate the electroporation efficiency by observing the expression of GFP in the GFP control group under an inverted fluorescence microscope 16 to 24 hours after electroporation. Some cell death (20 to 50%, depending on the electroporation program used) is expected after electroporation. >50% of the healthy cells should show GFP expression (Figure 2).
v. 16 to 24 hours after electroporation, change culture media to media with penicillin-streptomycin.
w. Continue to culture and passage the cells until the cells in the GFP control group have died off (usually within 1 to 2 weeks). Cells from the *Tp53* CRISPR group should continue to proliferate, which indicates successful immortalization.
x. Freeze down the established iMEFs and/or culture further for experiments.

| Pulse voltage (V) | Pulse width (ms) | Number of pulses |
| --- | --- | --- |
| 1650 | 20 | 1 |

**Figure 2:**
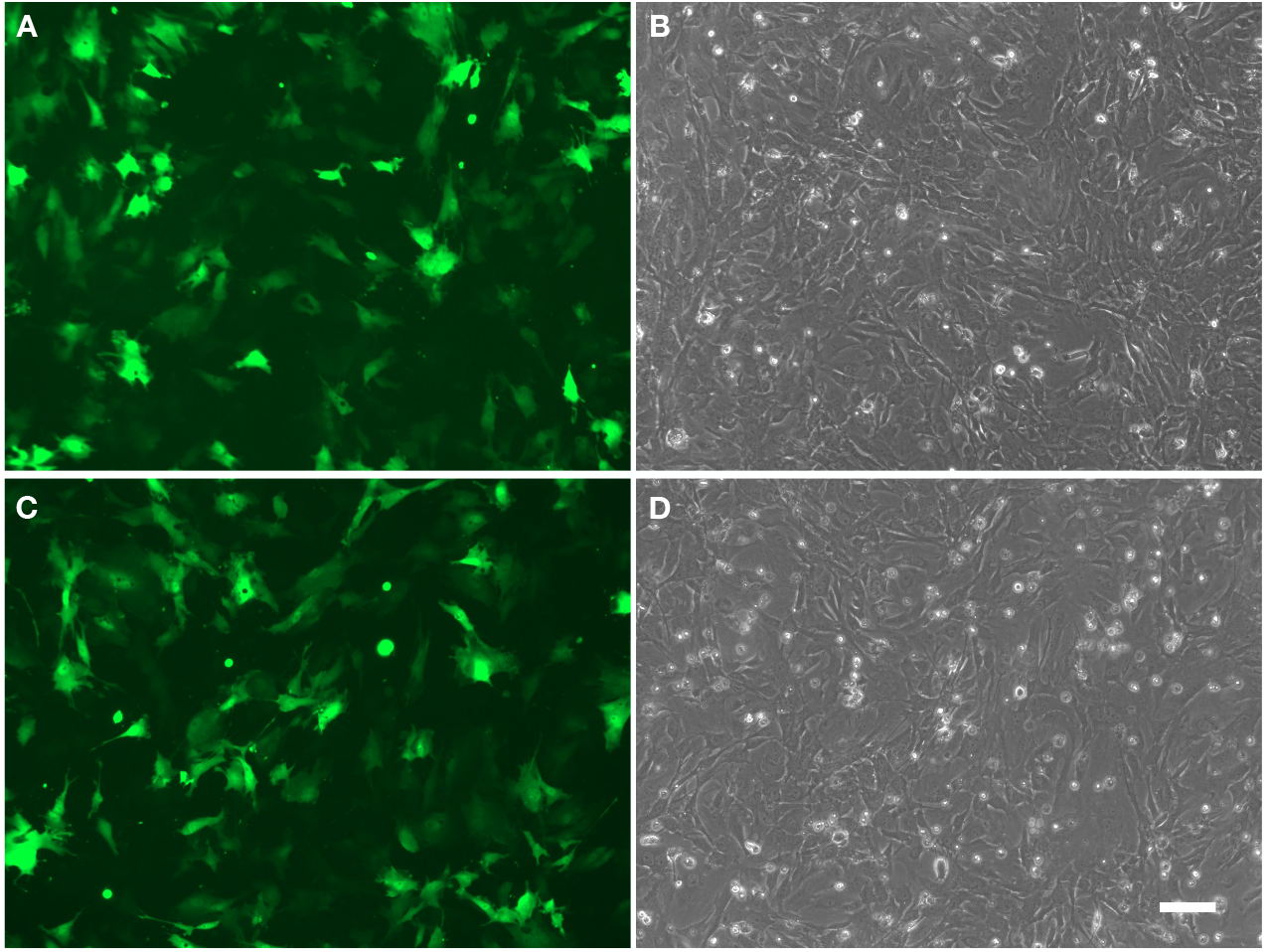
Evaluation of electroporation efficiency: Representative GFP (A, C) and phase contrast (B, D) images of MEFs one day post electroporation using program A (A, B) and program B (C, D). Scale bar: 100μm. Magnification: 10X.

###### Validation of Protocol

We have successfully used this protocol to generate more than 15 independent iMEF lines, including that described in Griffiths *et al, eLife*, 2024 (Figures 3 to 5, and the associated supplementary data figures) (Griffiths et al., 2024).

### Troubleshooting

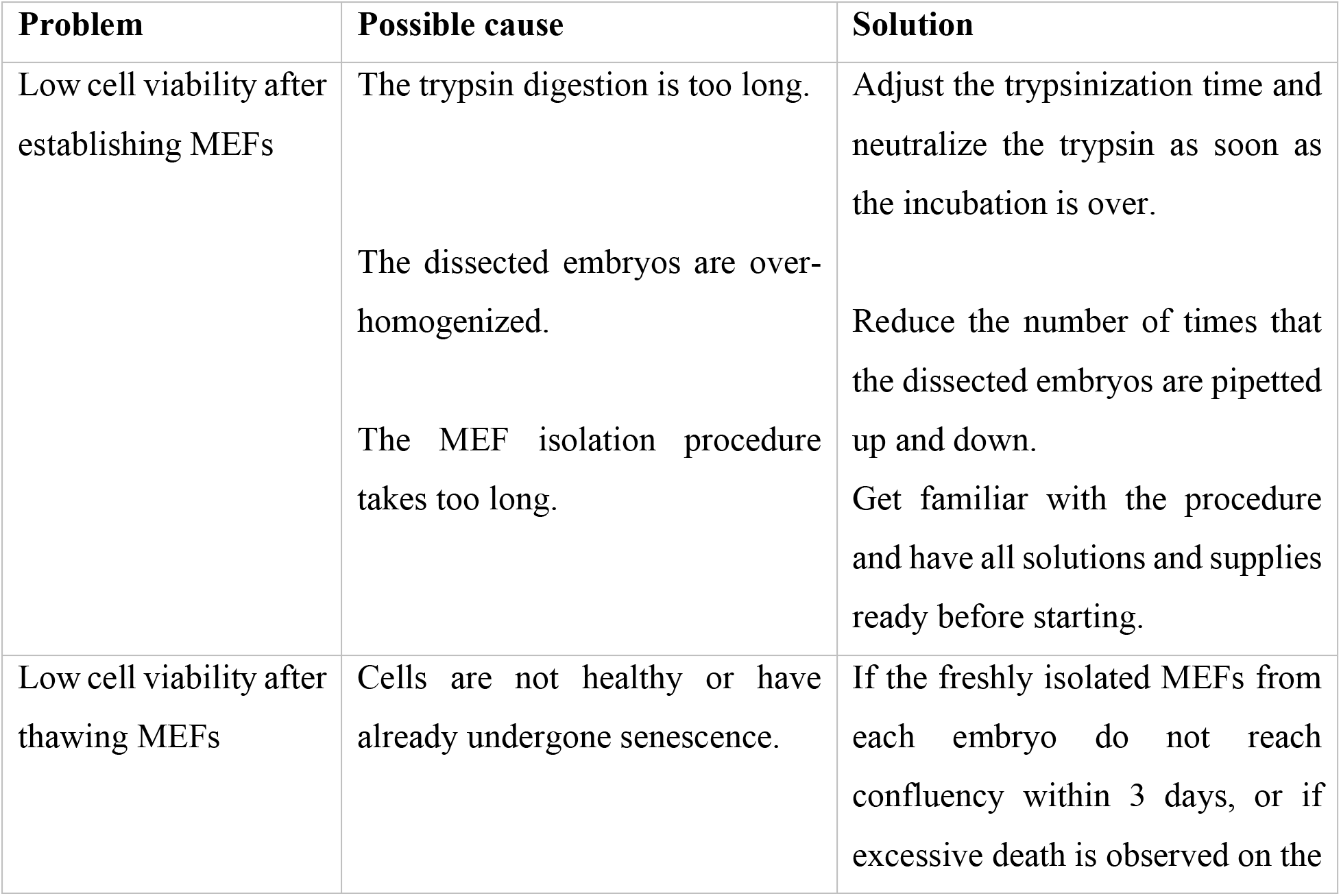
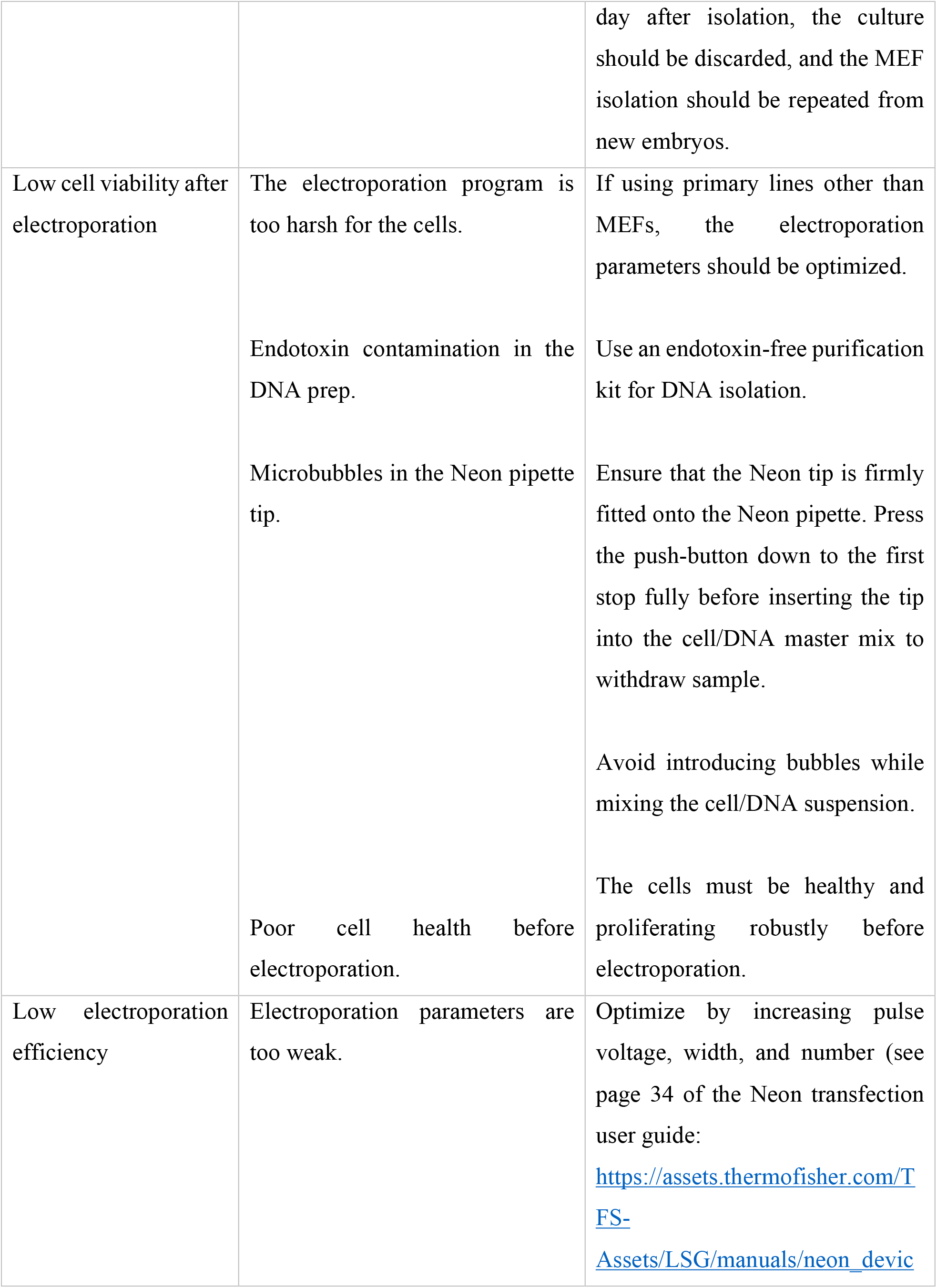
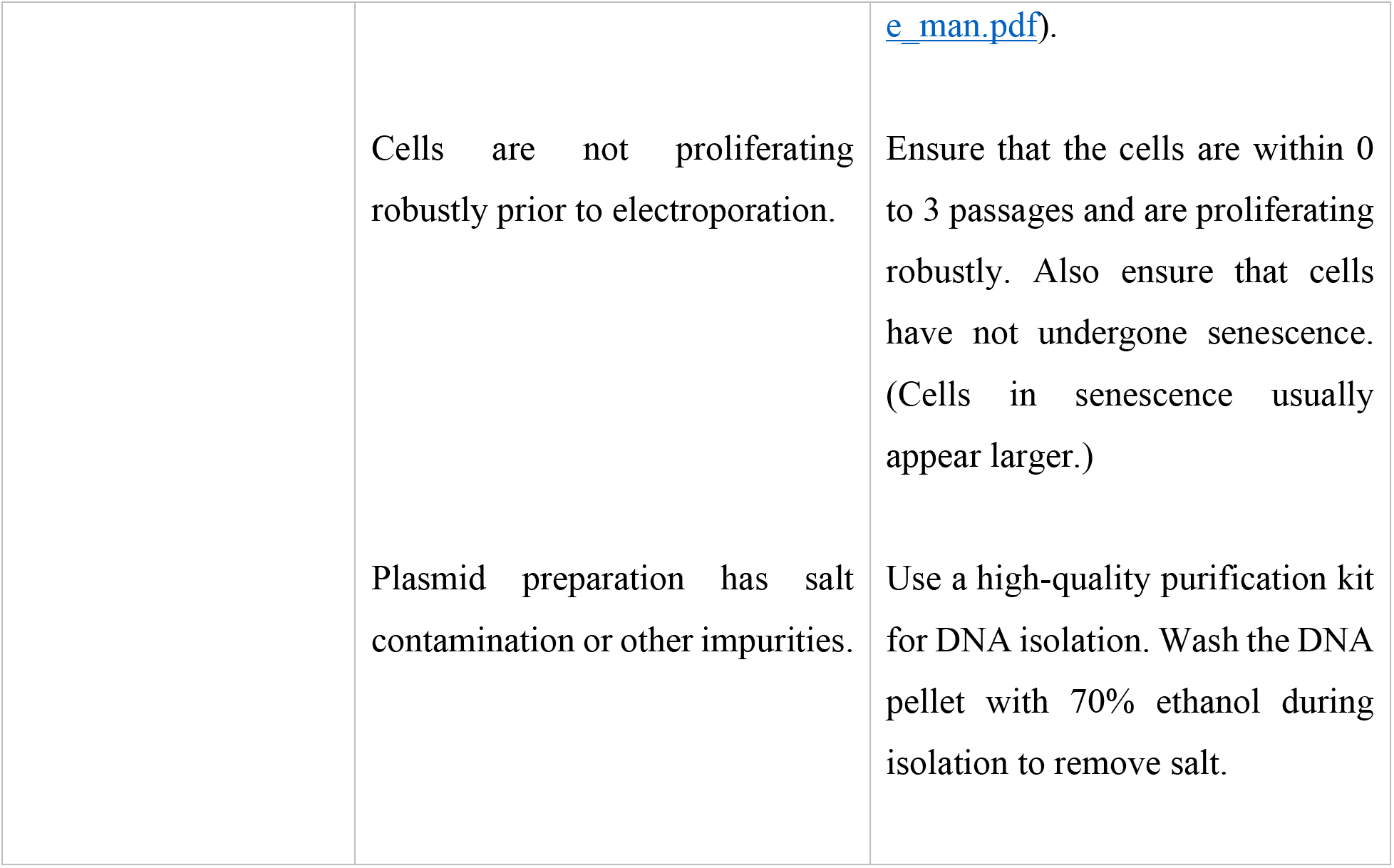

## Acknowledgments

This study was supported by the National Institutes of Health (1R35GM119574, 1R35GM144341) to H.H.H. This protocol was established to generate the iMEFs used in the following publication: Griffiths et al. (2024)

## Ethical considerations

1. All protocols using mice have been approved by Institutional Animal Care and Use Committee (IACUC), University of California, Davis.
2. All protocols involving DNA technology and biohazardous materials have been approved by the Institutional Biosafety Committee (IBC) at University of California, Davis.

